# Electric fields determine carbapenemase activity in class A β-lactamases

**DOI:** 10.1101/2023.11.04.565607

**Authors:** Hira Jabeen, Michael Beer, James Spencer, Marc W. van der Kamp, H. Adrian Bunzel, Adrian J. Mulholland

## Abstract

Antimicrobial resistance is a public health crisis. Limited understanding of the catalytic drivers in resistance-mediating enzymes such as β-lactamases hinders our ability to combat this crisis. Here, we dissect the catalytic contributions of active-site electric fields in class A β-lactamases. We studied the enzymatic hydrolysis of a carbapenem antibiotic by QM/MM molecular dynamics simulations and quantified active-site fields with a custom-made script. We discovered that the fields correlate well with activity and identified seven positions, some distal, that distinguish efficient carbapenemases. Electric-field analysis may help predict the activity of β-lactamases and guide antibiotic and enzyme design.

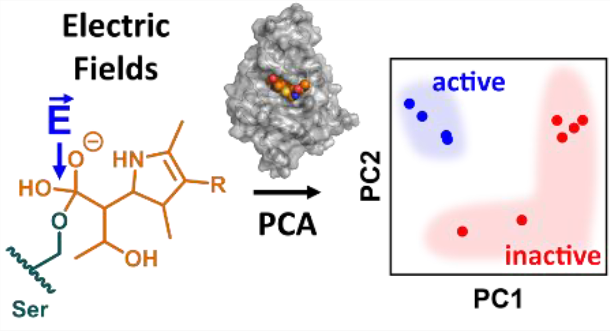

Electric field script: www.github.com/bunzela/FieldTools

---

Antimicrobial resistance is an escalating global crisis threatening human lives and many aspects of modern medicine. Around 1.2 million deaths annually are a direct result of infections with resistant pathogens.1 The overuse of antibiotics compounds this crisis and accelerates the evolution of antimicrobial resistance.^2,3^ Alarmingly, resistance development is outpacing the discovery of new antibiotics.^4,5^ Thus, as sequence information becomes more widely available as part of clinical microbiology workflows, there is an urgent need for reliable tools to predict the activity spectrum of emerging resistance genes and guide the design of next-generation antibiotics.^2^

In Gram-negative bacteria such as *Escherichia coli*, resistance to β-lactams, the most commonly prescribed antibiotic class, arises primarily through antibiotic hydrolysis by β-lactamases.^6–8^ In class A β-lactamases, hydrolysis follows a two-step mechanism (Fig. S1a). β-lactam breakdown commences with a nucleophilic attack of a catalytic serine on the amide carbonyl, leading to the formation of a covalent acyl-enzyme complex (AE) *via* a tetrahedral intermediate. Subsequently, these enzymes use a glutamate base that deprotonates a deacylating water to hydrolyze the AE complex *via* a second tetrahedral intermediate (TI, Fig. 1). Hydrolysis of the AE intermediate is slower than its formation for many β-lactams.^9,10^ Thus, various β-lactams, such as the carbapenem meropenem, have been engineered that slow down TI hydrolysis to effectively inhibit β-lactamase activity and prevent resistance.^11–13^

**Fig. 1.**
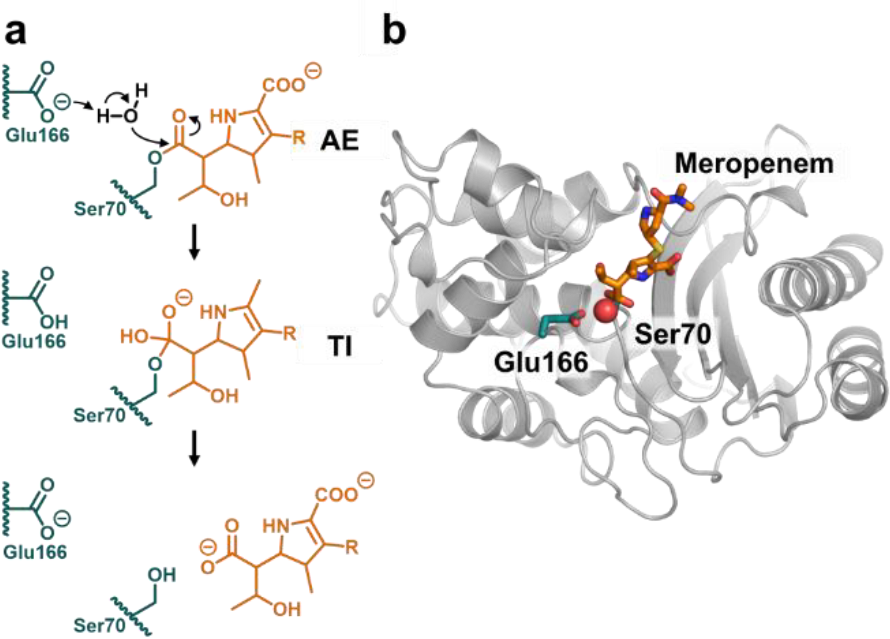
Carbapenem deacylation in class A β-lactamases. **(a)**To hydrolyze the acyl-enzyme intermediate (AE) *via* a tetrahedral intermediate (TI), class A β-lactamases use a carboxylate base (Glu166) that activates a deacylating water for nucleophilic attack on the acylated Ser70.^6^ **(b)** Model of the TEM-116:meropenem acyl-enzyme. (based on PDB ID: 1BT5;^16^ Glu, Ser: teal, meropenem: orange).

Nevertheless, clinically relevant class A β-lactamases, e.g., *Klebsiella pneumoniae* carbapenemase (KPC) have emerged that efficiently break down meropenem to confer resistance.^13–15^ Understanding the molecular origins of efficient AE hydrolysis, and how hydrolysis differs between enzymes and variants, is of crucial importance to elucidate β-lactamase activity and its relation to resistance phenotype.

Electrostatic interactions play a pivotal role in enzyme catalysis,^17–19^ and electrostatic stabilization of transient oxyanionic species such as the β-lactamase deacylation TI is key to catalysis in many hydrolytic enzymes.^20–22^ We hypothesized that residues critical for β-lactamase activity could be readily identified from their electrostatic interactions with the TI oxyanion. Due to the long-range nature of electrostatic interactions, the identified residues might even involve positions far away from the active site. Thus, electric field analysis might allow to pinpoint catalytically relevant remote residues, which represents a key challenge both for understanding existing biocatalysts and designing novel enzymes.^23,24^

Electric fields, experimentally observable from vibrational spectroscopy, represent a physical metric that can be used to quantify electrostatic effects.^17–19^ Here, we set out to perform atomistic molecular dynamics (MD) simulations to compare electric field effects in four class A β-lactamases with high hydrolytic activity towards the carbapenem meropenem (carbapenemases: KPC-2, NMC-A, SFC-1, SME-1) with those in six enzymes with insufficient activity to confer resistance (non-carbapenemases: BlaC, CTX-M-16, SHV-1, TEM-1, TEM-52, TEM-116).^11–16,25–32^ To that end, AE deacylation was simulated by hybrid quantum mechanics/molecular mechanics (QM/MM) MD using DFTB2/ff14SB.^9,33–38^ As benchmarked in our previous work,^9,34^ 2D umbrella sampling was used to simulate AE hydrolysis by following the deprotonation of the deacylating water and its nucleophilic attack upon the acyl-enzyme carbonyl. In addition, deacylation was simulated using the adaptive string method^35^, which increases sampling efficiency by projecting the collective variables onto a single reaction coordinate.

Both QM/MM sampling approaches give barriers (Δ*G*^‡^_calc_) that correlate well with experimental activity (Δ*G*^‡^_exp_; string method: *R*^2^ = 0.82; 2D umbrella sampling: *R*^2^ = 0.62; Fig. 2, S2 & S3). The calculated barriers are generally lower than the experimental activation energies (Tab. S1) because of limitations of the DFTB2 method, as noted previously.^9,34^ The string method gave lower barriers than those from 2D umbrella sampling. By its nature, the string method allows for better definition and more comprehensive sampling of the minimum free energy path. Sampling is further enhanced using replica exchange between windows in the adaptive string method implementation,^35^ but was not performed during 2D umbrella sampling. These factors likely decreased the calculated barriers. Nevertheless, the excellent correlation of Δ*G*^‡^_calc_ from either method with Δ*G*^‡^_exp_ suggests that the MD trajectories from both approaches are suitable to assess electrostatic effects promoting AE hydrolysis.

**Fig. 2.**
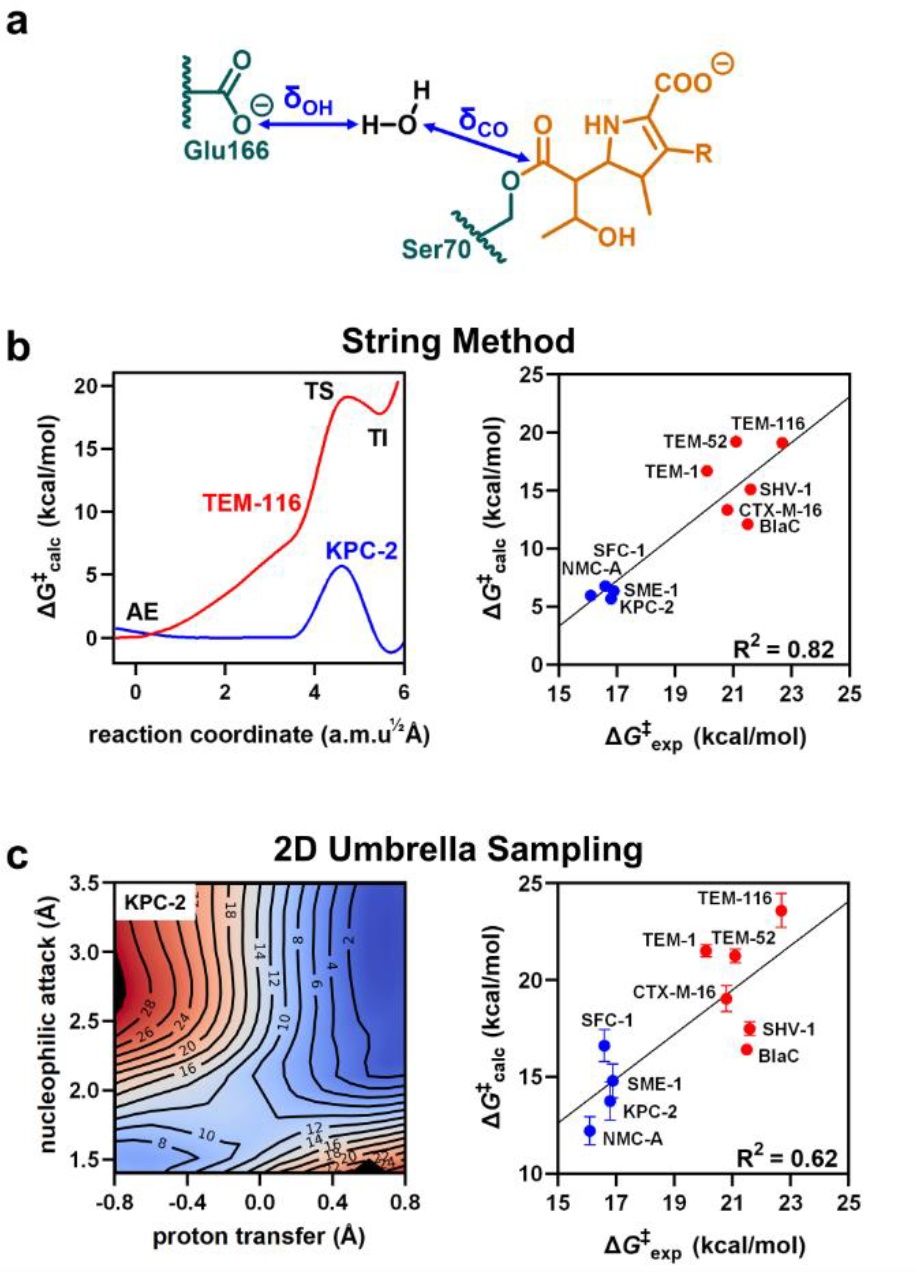
Calculated free energy barriers for AE deacylation agree with the experimental *k*cat values. **(a)** Reaction coordinates for AE hydrolysis comprise the proton transfer (δ_OH_) and nucleophilic attack (δ_CO_). **(b+c)** Δ*G*^‡^_calc_ from the String method **(b)** and 2D umbrella sampling **(c)** correlate well with Δ*G*‡^exp^ calculated based on *k*_cat_ (blue: carbapenemases; red: non-carbapenemases; Error bars represent the standard error of 10 independent calculations).

To investigate the electrostatic stabilization of the negative charge accumulating on the TI carbonyl oxygen, we determined electric fields along the β-lactam C=O bond. Fields in the AE, TS, and TI states were determined from the obtained QM/MM trajectories. Electric field vectors 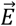 were first calculated at the C=O bond (Eq. S1) and subsequently projected onto the bond (Eq. S2) to give rise to the effective field (*E*_eff_) stabilizing the C=O dipole (Fig. 3a & b). Scripts for electric field calculation are available at www.github.com/bunzela/FieldTools. To mask effects intrinsic to the reaction, the ‘reactive’ part of the substrate, comprising the C=O bond with its adjacent carbon atoms and the deacylating water, was excluded from the *E*_eff_ calculations. Finally, we note that, while we focus on the results from the string method, results from 2D umbrella sampling trajectory data are qualitatively similar underscoring the significance of our findings (Fig. S3, S4 & S7).

**Fig. 3.**
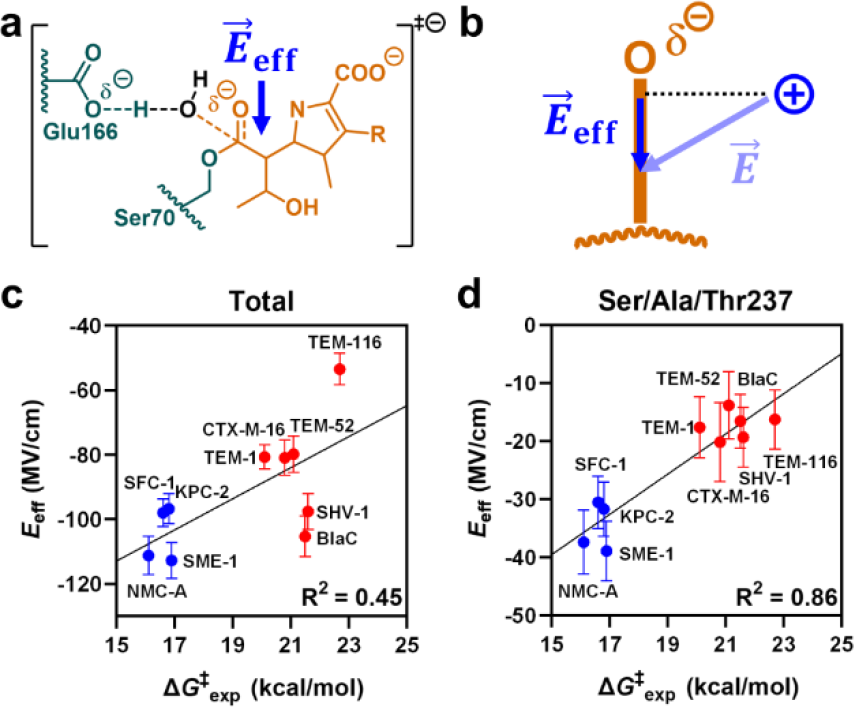
Electric fields in carbapenemases and non-carbapenemases. **(a+b)** *E*eff was calculated in the TS from the electric field 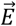projected by the enzyme along the carbonyl bond of the acylated carbapenem (orange). **(c+d)** While the total *E*_eff_ in the TS ensemble only weakly correlates with activity, the *E*_eff_ contributions of individual positions that interact with the oxyanion show good correlation. (*E*_eff_ values based on the string method; see Fig S4 for *E*_eff_ values based on 2D umbrella sampling and other per-residue values; error bars reflect the standard error of ten independent replicas).

The total *E*_eff_ was always negative and increased in magnitude during the reaction, resulting in better oxyanion stabilization in the TS and TI ensembles (Fig. 3 & S4, Tab. S2). Overall, *E*_eff_ values ranged between - 53 and -115 MV/cm in the TS ensemble, which agrees well with the fields observed in other enzymes that utilize oxyanion stabilization for catalysis.^18–22^ Although the magnitude of *E*_eff_ was overall marginally lower in the TS than the TI ensemble (Fig. S4), *E*_eff_ varied slightly more with Δ*G*^‡^_exp_ in the TS (4.8 (MV/cm)/(kcal/mol) compared to the TI (4.3 (MV/cm)/(kcal/mol)). Notably, the electric field in the AE changed less with activity compared to the TI and TS (1.6 (MV/cm)/(kcal/mol)), demonstrating the importance of reaction simulations for this purpose. The TI is often treated as a TS analog as it is more defined – and thus experimentally and computationally more accessible – than the TS. Although the two states are similar, our analysis also suggests that the TS is the more relevant state for studying effects on the reaction. Overall, our results show the importance of analyzing unstable states such as the TS to understand catalysis.

*E*_eff_ analysis overall reproduced the expected electrostatic oxyanion stabilization vital to β-lactamase activity. Nonetheless, *E*_eff_ reflects only one of several catalytic contributions, which might explain outliers observed in Fig. 3 and Fig. S4. For example, the non-carbapenemase BlaC contains the Asn132Gly substitution compared to the other carbapenemases studied here, which precludes a hydrogen bond with the carbapenem 6α-hydroxyethyl group and decreases substrate preorganization (Fig. S5b).^6,11^ Other factors previously shown to impact carbapenem hydrolysis include control of water-mediated substrate tautomerization, organization of the active site through extended hydrogen-bonding networks, and a disulfide bond between Cys69 and Cys238.^10,27,39–42^ We stress that *E*_eff_ and Δ*G*^‡^_exp_ showed the expected trends when analyzing the three studied TEM variants, which differ by no more than five mutations. These data suggest that electric field analyses can identify small but catalytically important electrostatic effects resulting from limited sequence variation in closely related enzyme systems.

To identify the residues that provide the major contributions to the electrostatic effect, the total *E*_eff_ can be readily partitioned into per-residue contributions. To that end, we performed a structure-based sequence alignment of the studied β-lactamases. Subsequently, *E*_eff_ values were calculated for each residue, apart from those in loops that vary in length between the enzymes (residues 26-28, 50-56, 85-90, 141-144, 240-241, 268-274, Fig. S6). For reference, our analysis will thus use the residue numbers of the TEM variants. Strong correlation of the residue-specific *E*_eff_ with Δ*G*^‡^_exp_ pinpoints several residues, of which some are known to be directly involved in electrostatic oxyanion stabilization, such as the oxyanion-hole donating residue 237 (Fig. 3C & S7).

The calculated per-residue *E*_eff_ values were subjected to principal component analysis (PCA) to identify sites that are the primary contributors to electric field differences between the enzymes. PCA is a statistical method that can reveal trends in complex datasets by projecting data onto a smaller set of principal components. The landscape of the first and second principal components (PC1, PC2) revealed a defined cluster for the carbapenemases, while non-carbapenemases populated a broad but distinct distribution. Thus, PCA of the per-residue *E*eff data clearly distinguishes carbapenemases from non-carbapenemases (Fig. 4a).

**Fig. 4.**
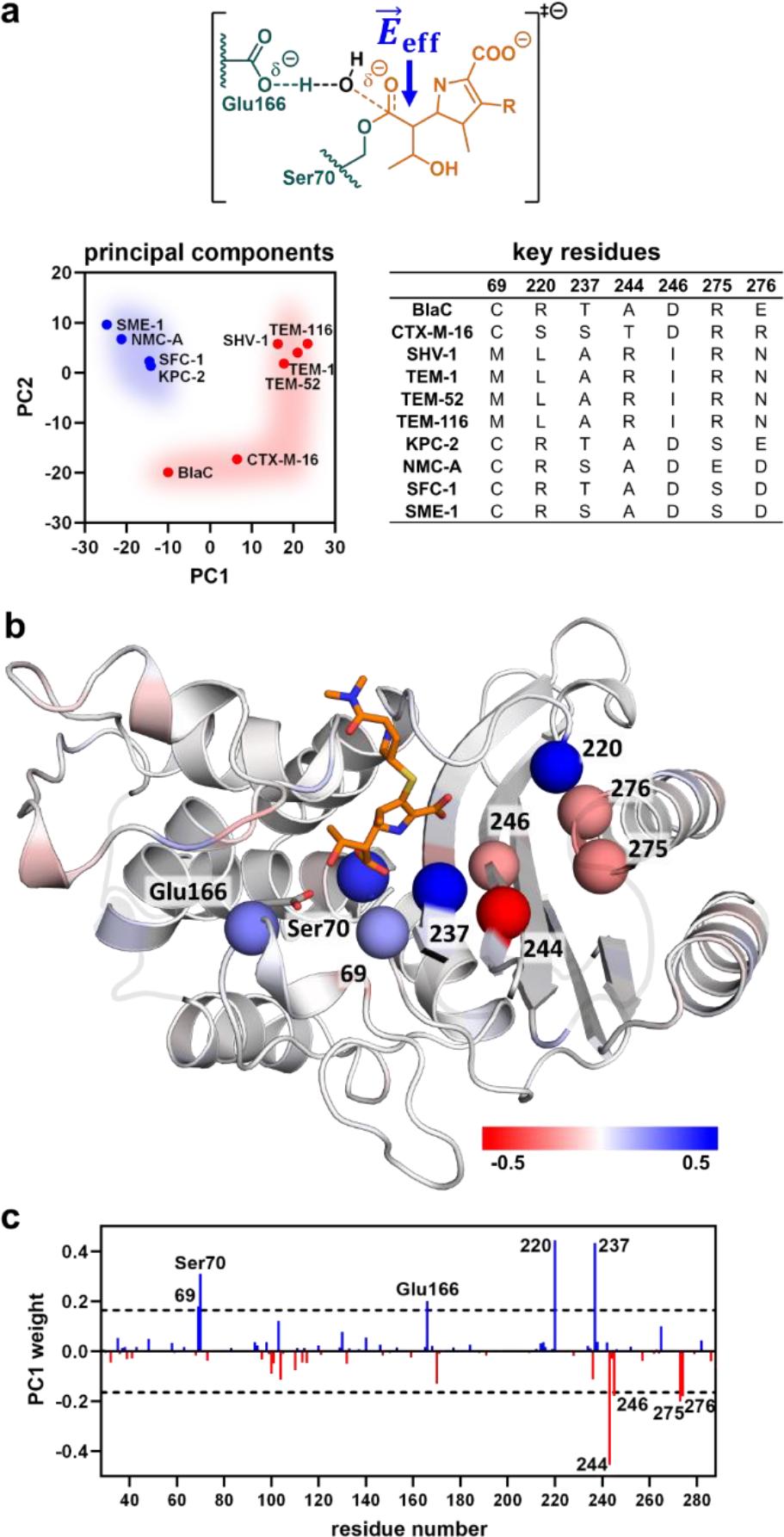
Contribution of individual residues to the electric field. **(a)** PCA of the per-residue *E*_eff_ clearly distinguishes carbapenemases (blue) from non-carbapenemases (red, shaded areas added for illustration only). **(b+c)** Principal component 1 (PC1) weights reveal that a cluster of seven residues and the catalytic residues (spheres) dominates *E*_eff_ (blue: beneficial; red: detrimental, meropenem: orange; dashed line: employed cutoff; PCA based on string method; see Fig. S7 for PCA based on 2D umbrella sampling).

Principal component weights can provide detailed insights into which parameter (here: which position in the structure) has the strongest effect. For the first principal component (PC1), weights of the two conserved catalytic residues and seven positions, some distal, differed by ≥2.5 standard deviations from the average weight (Figure 4b & c). Residues with a significant positive weight comprise the catalytic base Glu166, the acylated Ser70, and residue 237, whose backbone amide constitutes the oxyanion hole together with Ser70. Two additional residues, 69 and 220, that also cluster around the oxyanion hole had positive weights. Residues 244, 246, 275, and 276, which likewise clustered around the substrate oxyanion, had significant negative weights. Comparison of the per-residue weights with their correlation of *E*_eff_ with Δ*G*^‡^_exp_ shows that positive weights correspond to residues that exert catalytically more beneficial fields as activity increases. In contrast, negative weights indicate residues whose *E*_eff_ values are worse in the more active enzymes (Fig. S4 & S7).

Intriguingly, both the PCA and the per-residue correlations of *E*_eff_ with Δ*G*^‡^_exp_ identified similar residues with substantial electrostatic contributions (Fig. S4 & S7). We emphasize that neither Δ*G*^‡^_calc_ nor Δ*G*^‡^_exp_ were included in the PCA. The PCA is thus not biased by any activity data and solely reflects changes in the electrostatics between variants. Consequently, the total electric field effect in the studied β-lactamases is dominated by the catalytic residues and a cluster of seven residues around the oxyanion hole (Fig. 4b), with PCA reliably pinpointing these relevant, and partially non-obvious, residues.

All carbapenemases studied here have an arginine residue at position 220, while the non-carbapenemases all have at least one arginine at positions 244, 275, or 276 (Fig. S1b+c). Notably, the only non-carbapenemase with Arg220 is BlaC, which also shows an unexpectedly strong overall *E*_eff_ as discussed before (Fig. 3c). Class A β-lactamase activity is demonstrably affected by the presence of positively charged arginine residues in the vicinity of the oxyanion hole, but understanding the differential effects of a charged residue at varying positions has been challenging.^43–48^ For instance, the R244A mutation in TEM-1 impairs hydrolysis of various penicillin and cephalosporin-based β-lactams, but introduction of arginines at positions 220, 272, or 276 can partially recover activity.^46^ Our electric field calculations likewise confirm that arginine residues near the oxygen side of the substrate C-O bond generally boost electrostatic catalysis. In addition, our work provides a molecular description of the differential effect of arginine at various positions. Per-residue *E*_eff_ values show that an arginine at position 220 is catalytically superior for meropenem hydrolysis compared to positions 244, 275, or 276. (Fig. S7). Our analysis suggests that the differential *E*_eff_ of arginine at various positions is probably an important discriminator for carbapenemase activity.

In summary, combining electric field calculations with principal component analysis enabled a detailed per-residue study of the electrostatic determinants of β-lactamase activity. Our analysis revealed that a cluster of seven residues dominates electrostatic oxyanion stabilization and, together with the two established catalytic residues, differentiates carbapenemases from non-carbapenemases. Given that resistance genes in pathogenic strains from patient samples can now be reliably identified by genome sequencing,^2^ we anticipate that electric field calculations may aid in predicting the resistance spectrum of emerging enzymes. In that regard, our assay could potentially be accelerated by simulation of the metastable TI state alone to be time-compatible with rapid sequencing and inform clinical guidance.

Our work underscores the potential of integrating reaction simulations and electric field calculations to discern effects along the reaction coordinate.^49–54^ Simulation offers various advantages compared to experimental methods that typically rely on vibrational probes: (1) computational assays are not limited to transition state analogs or ground state substrates; (2) specific states along the catalytic cycle can readily be studied; (3) electric fields can easily be partitioned into individual components

Dissecting per-residue electrostatic effects by PCA was instrumental in pinpointing residues key to determining carbapenemase activity. Notably, PCA allowed us to dissect electrostatic effects without biasing the analysis by comparing to reaction barriers. Nonetheless, the analysis reliably reproduced the catalytically important residues found by comparing *E*_eff_ with Δ*G*^‡^_exp_. We note that using residue-based principal component weights to identify contributions to catalysis is likely not limited to electric field effects and could potentially be used to dissect any residue-based phenomenon. Finally, total or residue-based *E*_eff_ values could be used as features in machine learning models aimed at understanding and designing catalytic activity.^55,56^

In conclusion, our work shows that highly optimized electric fields in naturally evolved β-lactamases can give rise to specific antibiotic resistance phenotypes. The tools developed here not only shed light on the molecular origins of electrostatic catalysis in class A β-lactamases, but also pave the way for dissecting electrostatic effects across a wide range of enzymes. Our FieldTools script allows the user to rapidly assess electric fields and is thus amenable to computational design (www.github.com/bunzela/FieldTools). Targeting electric fields by design might pave the way to create enzymes more efficiently,^57–60^ and reliably introduce remote catalytic interactions.^23,24^ By demonstrating how nature evolves to evade antibiotic efficacy, our work enhances our ability to predict the emergence of antimicrobial resistance and provides a strategic framework for the development of next-generation antibiotics.

## AUTHOR INFORMATION

We encourage critical feedback and suggestions on our preprint. Please submit any comments to.

## AUTHOR CONTRIBUTIONS

HJ, HAB, and AJM conceived the experiments. HJ performed the simulations and analysis with support from HAB. MB, MWvdK, and JS contributed to the data interpretation. HJ and HAB wrote the manuscript with input from all authors.

## ACKNOWLEDGEMENTS

This work was conducted using the computational facilities of the Advanced Computing Research Centre, University of Bristol. We thank Kirill Zinovjev for helping with setting up the adaptive string method. We thank Christopher Fröhlich for feedback on our manuscript.

## DATA AND CODE AVAILABILITY

The FieldTools electric field calculation script is available at https://github.com/bunzela/FieldTools. For questions regarding the FieldTools script, we strongly encourage contacting HAB directly.

