## Supplementary Information for "Electric fields determine carbapenemase activity in class A β-lactamases"

### Supporting information for:
### Electric fields determine carbapenemase activity in class A β-lactamases

Hira Jabeen,[a] Michael Beer,[b] James Spencer,[b] Marc W. van der Kamp,[a,d] H. Adrian Bunzel,[a,c,*] and Adrian J. Mulholland[a,*]

[a] *Centre for Computational Chemistry, School of Chemistry, University of Bristol, UK*
[b] *School of Cellular and Molecular Medicine, University of Bristol, UK*
[c] *Department of Biosystem Science and Engineering, ETH Zurich, CH*
[d] *School of Biochemistry, University of Bristol, UK*

#### 2. Methods

MD simulations were based on previously built systems and generated parameters (see 2.1 System setup), and the 2D Umbrella sampling (see 2.3 2D umbrella sampling free energy calculations) were run similarly to tested and benchmarked methods.[9,34] Where simulations were available,[9] starting structures were taken from 300 ps QM/MM simulations from this previous work.

##### 2.1. System setup

We studied ten class A β-lactamases selected for their clinical relevance and capacity to either hydrolyze carbapenems (KPC-2, NMC-A, SFC-1, and SME-1), or that were inhibited by carbapenems (BlaC, CTX-M-16, SHV-1, TEM-1, TEM-52, and TEM-116).[11–15] [9,34]Acyl-enzyme (AE) models were based on X-ray crystal structures with the following PDB IDs: 3DWZ, 1YLW, 2ZD8, 1M40, 1HTZ, 1BT5, 2OV5, 1BUE, 4EV4, and 1DY6.[14,16,25–32] For BlaC and SHV-1, meropenem-bound acyl-enzyme crystal structures were available. The TEM-1 crystal structure was obtained in complex with a boronic acid transition state analog, which was replaced by meropenem using. The TEM-116 crystal structure was obtained in complex with imipenem, which was replaced by meropenem. The SFC-1 crystal structure contained a mutation (E166A), which was reverted to wild type based on an S70A meropenem-bound variant (PDB: 4EUZ). The apoenzyme structures of CTX-M-16, TEM-52, KPC-2, NMC-A, and SME-1 were aligned with the SFC-1-meropenem complex to build the respective acyl-enzyme using the meropenem coordinates from the SFC-1 structure.

The AE complex with meropenem in its $\Delta^2$ tautomer was parametrized with RESP charges based on HF/6-31+G(d) density and GAFF parameters (https://doi.org/10.6084/m9.figshare.8158097.v1). The protein was parametrized with the ff12SB forcefield. Apart from the deacylating water, all crystallographic water molecules were deleted. Propka3.1[36] was used to calculate the protonation state, and hydrogens were added using tLeap.[37] Each system was solvated with TIP4P-Ew in a 10 Å box and neutralized by adding chloride or sodium ions. The QM region was treated at the DFTB2 level. The QM region comprised 41 atoms and 3 link atoms: covalently bound meropenem, from the CB of Ser70 to the S of meropenem, the sidechain of Glu166 from its CG atom, and the deacylating water.

Restraints were added to the simulations to avoid large conformational changes and unwanted bond-breaking in the QM region.[9,34] A weak harmonic restraint (>2.5 Å, 10 kcal/mol/Å$^2$ force constant) was applied between the carbonyl oxygen and amine, which formed the β-lactam in the meropenem substrate, to maintain a productive conformation. Likewise, a harmonic restraint (>1.2 Å, 10 kcal/mol/Å$^2$ force constant) was applied to the carbonyl carbon and oxygen of the acylated serine to prevent bond breakage. Three additional restraints (>1.2 Å, 50 kcal/mol/Å$^2$ force constant) were applied on the water O-H, hydroxyethyl O-H, and β-lactam amine N-H bonds to prevent proton transfer.

#### 2.2. String method free energy calculations

The adaptive string method,[35] incorporated in the Amber18[37] Sander module, was used to calculate the minimum free energy path. The string method can effectively calculate minimal free energy path in multi-dimensional full energy surfaces by projecting multiple collective variables onto a single reaction coordinate.[35] Two collective variables were chosen to calculate free energy profiles (Fig. 2a). The proton transfer was captured by the distance between the hydrogen of the deacylating water and the carboxylate oxygen of Glu166 ($\delta_{OH}$). The distance between the deacylating water's oxygen and the meropenem carbonyl carbon (δCO) defined the nucleophilic attack. The string method requires the definition of the reaction's start and end point for each collective variable. The initial guess file contained two points ([1.80645, 3.52351] and [1.00866, 1.46509]) describing reaction coordinates that are slightly before the AE or after the TI. These points were optimized to ensure that the distribution of each coordinate along the equilibrate string is not biased by the endpoints. The optimization was done by adjusting the endpoints until the equilibrated string would not overshoot the given points. String calculations were performed for 28 string nodes, using the same QM/MM equilibrated starting structure as input for each node. String calculations were run for a total of 500 ps. The string was equilibrated for 25 ps plus an additional 25 ps as defined in the STOP_STRING file, and the free energy profile was subsequently obtained from sampling for 450 ps (Fig. S2).

#### 2.3. 2D umbrella sampling free energy calculations

2D umbrella sampling was used to calculate free energy surfaces and barriers. To explore the free-energy surface, single structures equilibrated through 300 ps QM/MM were submitted to 2D umbrella sampling. Reaction coordinates ranging from 3.5 to 1.4 for the nucleophilic attack and 0.8 to -0.8 for the deprotonation were sampled resulting in 374 windows. This range was extended compared to our previous work to ensure that the broad AE energy well was sampled exhaustively. Based on the equilibrated structure after 300 ps, umbrella sampling was done for 2 ps on each window to obtain 2D energy surfaces (Fig. 2c+S2, final 2D surface show the combined data from this 2 ps sampling and the 100 ps sampling of the minimum energy path described below). All studied β-lactamases have similar minimum free energy paths on full energy surfaces obtained through minimum energy path surface analysis (MEPSA).[38] The final potential of mean force for the two reaction coordinates was obtained through the weighted-histogram analysis method (WHAM).[33] Subsequently, free energy reaction barriers were calculated starting from ten independent configurations 15 ps apart from the 300 ps QM/MM simulations of the AE complex. For some of the picked frames, umbrella sampling did not run to completion due to the misalignment of the deacylating water at the active site in an inappropriate position for nucleophilic attack and deprotonation. If the umbrella sampling did not run properly, simulations were discarded and restarted with a different frame. 28 windows based on the minimum free energy path along the reaction coordinates were chosen, starting from the reactant structure towards the product structure for more extensive 100 ps umbrella sampling for each window resulting in the barriers in Fig. 2c.

#### 2.4. Electric field calculations

The electric field projected by the enzyme onto the reactive carbonyl moiety of the β-lactam was calculated using FieldTools (https://github.com/bunzela/FieldTools). FieldTools relies on Coulomb's law using the point charges from the system's topology file and the coordinates from the input trajectory to calculate electric fields along a target bond or at a atom. The electric field vector $\vec{E}$ was calculated from the Coulomb constant k, and the vector $\vec{r}_i$ from the carbonyl carbon (C=O) to all charges $Q_i$ in the system (Eq. 1). Subsequently, the effective field $E_{eff}$ projected along the carbonyl carbon was calculated by the scalar product of the directional unity vector $\vec{d}$ along the carbonyl carbon (C=O) and the total field $\vec{E}$ (Eq. 2).

$$\vec{E} = \sum_i k \frac{Q_i}{\vec{r}_i^2} \quad \text{Eq. S1}$$

$$E_{eff} = \vec{E} \cdot \vec{d} \quad \text{Eq. S2}$$

#### 3. Supporting Tables

##### Tab. S1 | Experimental and calculated activation energies.

| β-lactamase | $k_{cat}$ [a] ($s^{-1}$) | $\Delta^{‡}G_{exp}$ (kcal/mol) | $\Delta^{‡}G_{calc}$ US (kcal/mol) | $\Delta^{‡}G_{calc}$ String (kcal/mol) |
|---|---|---|---|---|
| BlaC | 0.0017 | 21.5 | 19.0 ± 1.9 | 11.9 |
| CTX-M-16 | 0.0042 | 20.8 | 18.4 ± 0.7 | 13.3 |
| SHV-1 | 0.0013 | 21.6 | 17.3 ± 0.4 | 15.0 |
| TEM-1 | 0.0625 | 21.1 | 21.5 ± 0.3 | 16.7 |
| TEM-52 | 0.0082 | 22.1 | 21.0 ± 0.4 | 19.1 |
| TEM-116 | 0.0023 | 22.7 | 23.6 ± 0.9 | 19.1 |
| KPC-2 | 3.6 | 16.8 | 13.0 ± 0.9 | 5.6 |
| NMC-A | 12.0 | 16.1 | 12.8 ± 0.5 | 6.8 |
| SFC-1 | 6.5 | 16.6 | 16.5 ± 0.5 | 4.0 |
| SME-1 | 3.2 | 16.9 | 14.0 ± 0.5 | 6.3 |

[a] Experimental $k_{cat}$ values resulting in $\Delta^{‡}G_{exp}$ were taken from refs. [11–15].

##### Tab. S2 | Calculated electric fields.

| β-lactamase | $\Delta^{‡}G_{exp}$ (kcal/mol) | Electric Field 2D Umbrella Sampling (MV/cm) | Electric Field String Method (MV/cm) |
|---|---|---|---|
| | | Total | Total |
| BlaC | 21.5 | -90 ± 8 | -105 ± 6 |
| CTX-M-16 | 20.8 | -88 ± 6 | -81 ± 5 |
| SHV-1 | 21.6 | -96 ± 2 | -98 ± 6 |
| TEM-1 | 20.1 | -70 ± 3 | -67 ± 4 |
| TEM-52 | 21.1 | -78 ± 11 | -80 ± 6 |
| TEM-116 | 22.7 | -72 ± 10 | -53 ± 5 |
| KPC-2 | 16.8 | -96 ± 4 | -97 ± 5 |
| NMC-A | 16.1 | -107 ± 4 | -111 ± 6 |
| SFC-1 | 16.6 | -100 ± 9 | -99± 4 |
| SME-1 | 16.9 | -101 ± 5 | -113 ± 6 |
| Slope ((MV/cm)/(kcal/mol)) [a] | | 3.6 | 5.02 |

[a] slopes were calculated in the transition state produced with QM/MM trajectories from either string or 2D umbrella sampling

#### 4. Supporting Figures

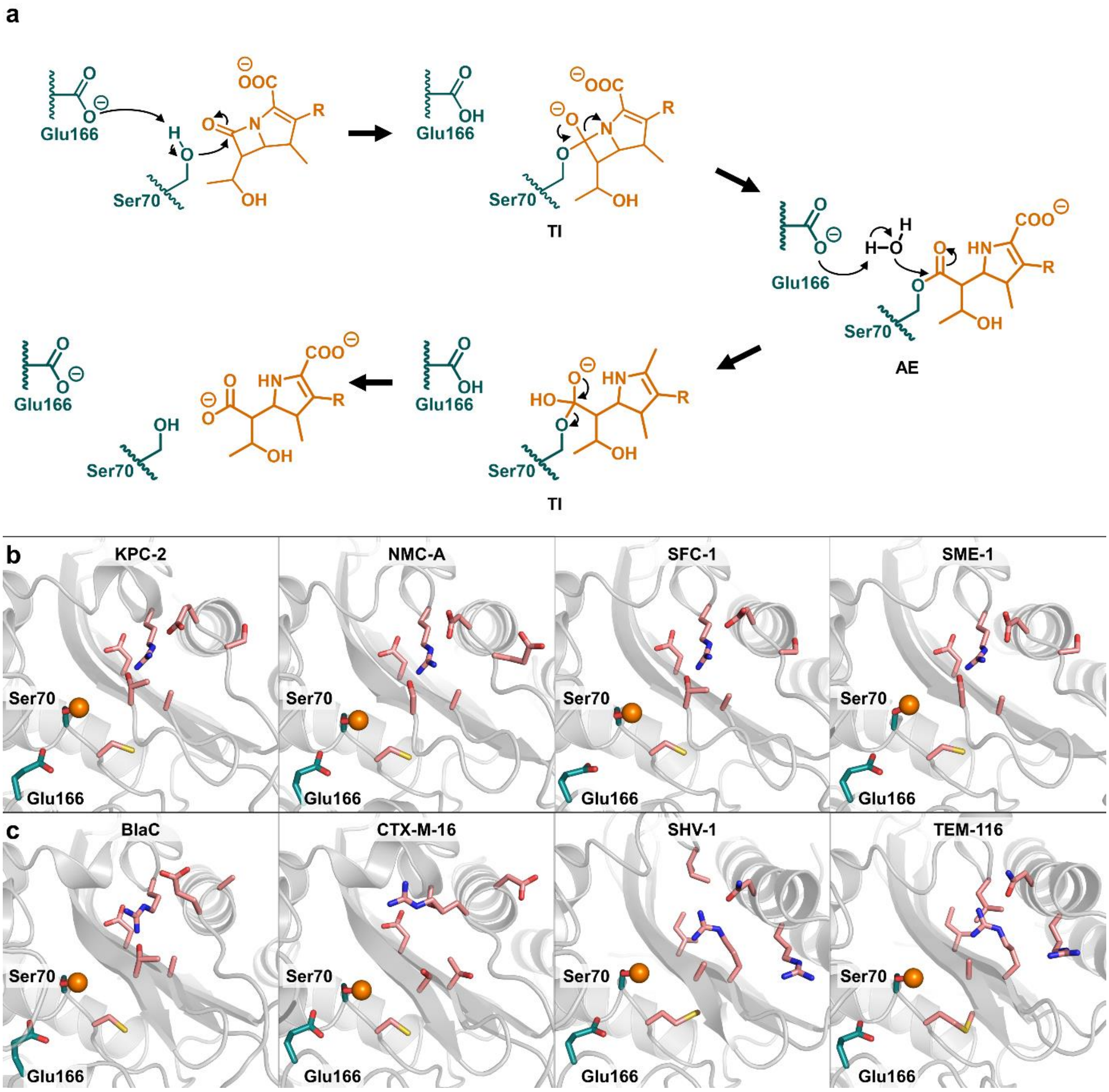

**Fig. S1 | Carbapenemase mechanism and active sites.**

**(a)** Class A β-lactamases cleave the carbapenem meropenem (orange) in a two-step mechanism. In the first step, an acyl-enzyme state (AE) is formed via a tetrahedral intermediate (TI). The AE is then hydrolyzed via a second tetrahedral intermediate (TI). The latter half of the reaction is rate-limiting in the studied carbapenemases.[11–15] **(b)** Active sites of the studied carbapenemases and **(c)** non-carbapenemases. The carbonyl carbon of meropenem is indicated as an orange sphere. The catalytic Ser70 and Glu166 are shown in teal and the other residues identified as catalytically important as shown in salmon. Because all studied TEM variants shared the same key residues, only TEM-116 is shown.

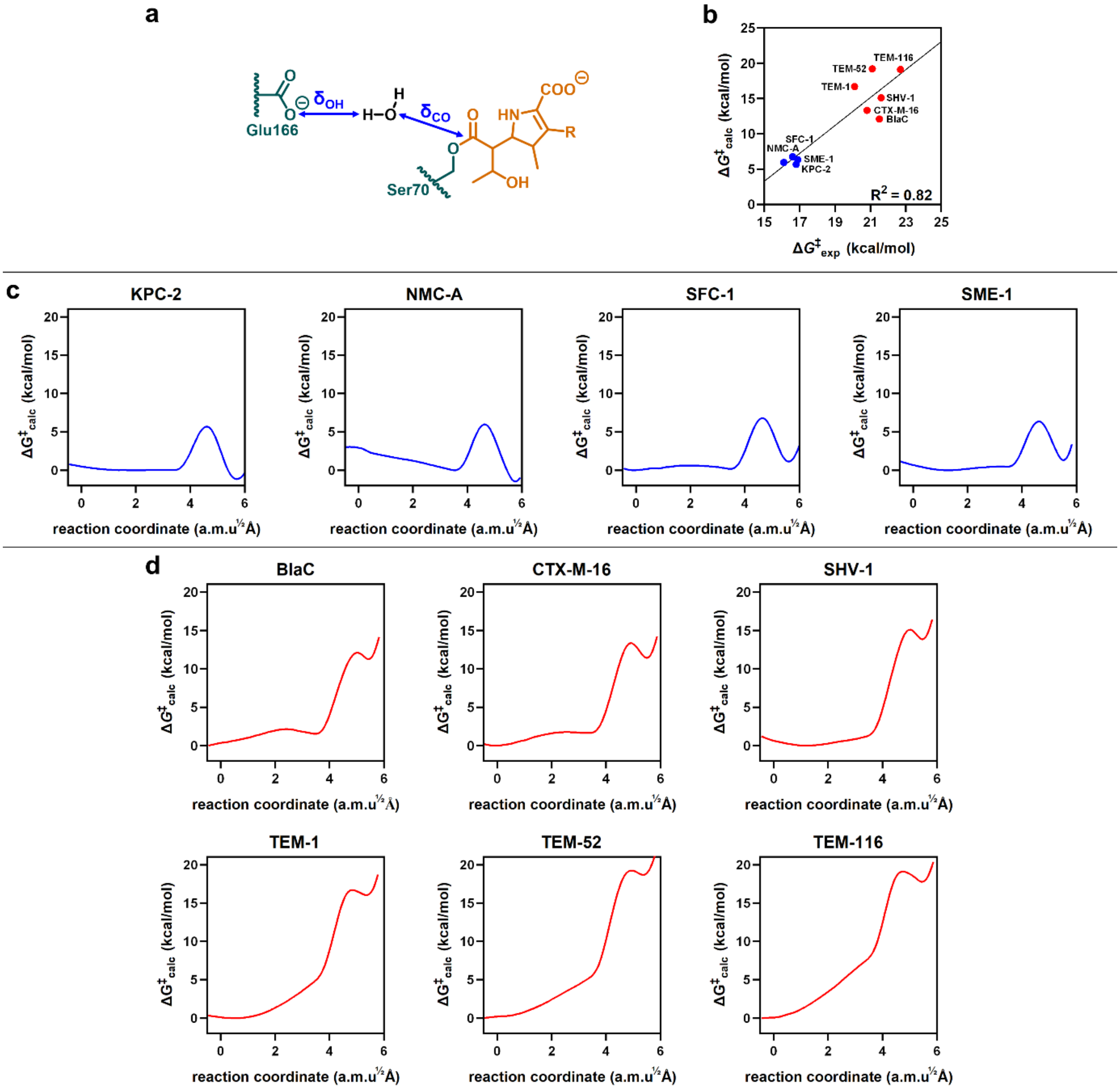

**Fig. S2 | Free energy barriers from string calculations.**

**(a)** The $\delta_{OH}$ and $\delta_{CO}$ distances were sampled during the string calculation. **(b)** Free energies calculated with the String method correlate well with the experimental activation energies (blue: carbapenemases; red: non-carbapenemases). **(c)** Potential mean force (PMF) profiles of carbapenemases measured along minimum free energy path. The highest energy point on each profile represents the transition state (TS) before tetrahedral intermediate (TI) formation. **(d)** PMF profiles of six carbapenem-inhibiting enzymes. Non-carbapenemases show higher deacylation barriers and later transition states than active carbapenemases.

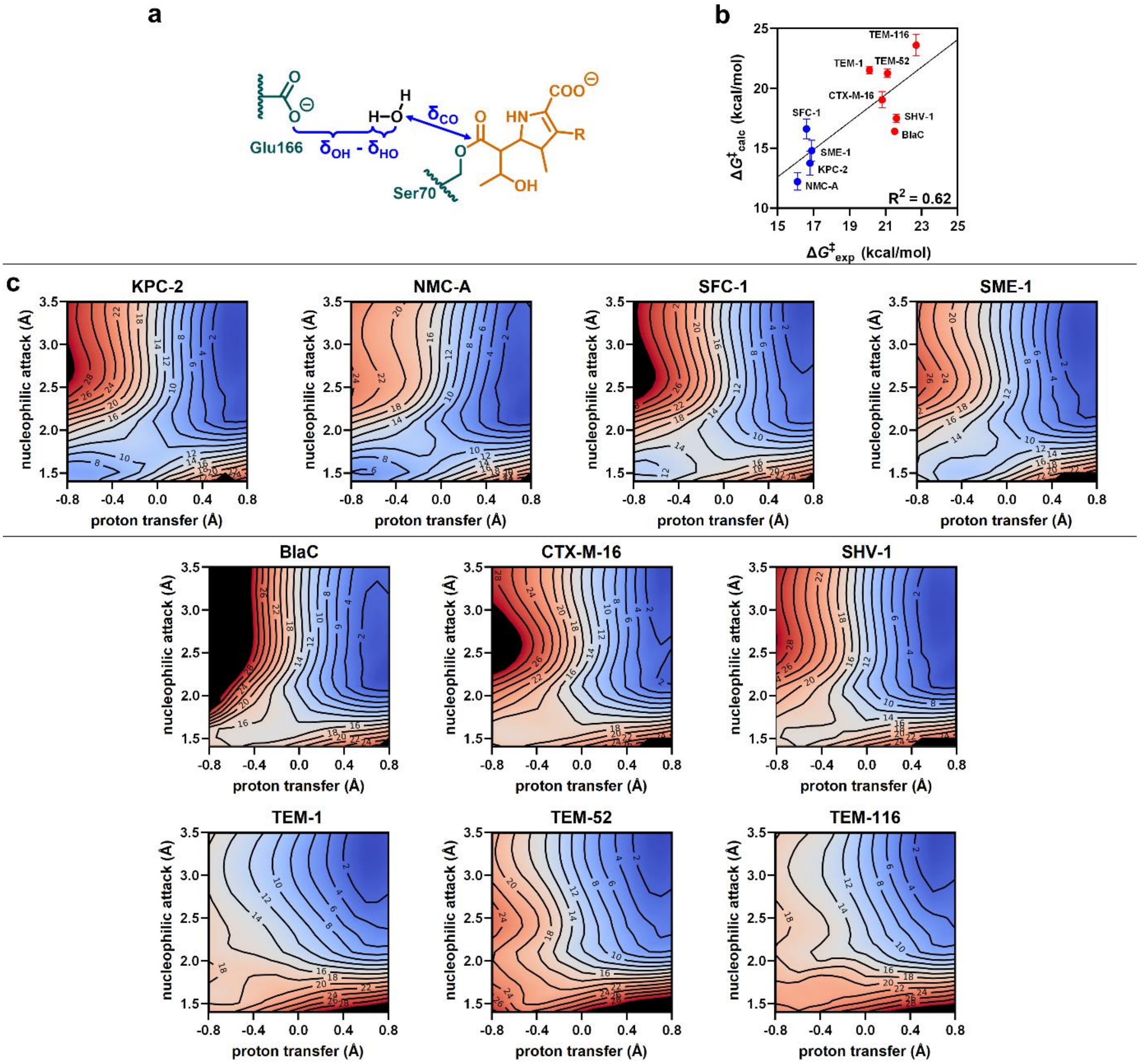

**Fig. S3 | Free energy barriers from 2D umbrella sampling.**

**(a)** The reaction coordinates sampled during 2D umbrella sampling comprised the proton transfer defined by the distance between the Glu166 carboxylate O and the deacylating water H minus the distance between the deacylating water H and O ($\delta_{OH}$ - $\delta_{HO}$), and the nucleophilic attack of water onto the AE intermediate ($\delta_{CO}$). **(b)** Free energy barriers calculated from 100 ps 2D umbrella sampling along the minimum free energy path correlate well with the experimental activation energies (blue: carbapenemases; red: non-carbapenemases). **(c)** 2D free energy surfaces of carbapenemases and **(d)** non-carbapenemases were calculated for the deacylation reaction for 2 ps sampling of each window (surfaces represent combined data from the 2 ps calculations and the 100 ps minimum free energy path sampling). Similar to the TS energies from the String calculations (Fig. S2), the difference in TS and TI energies is minor in the non-carbapenemases

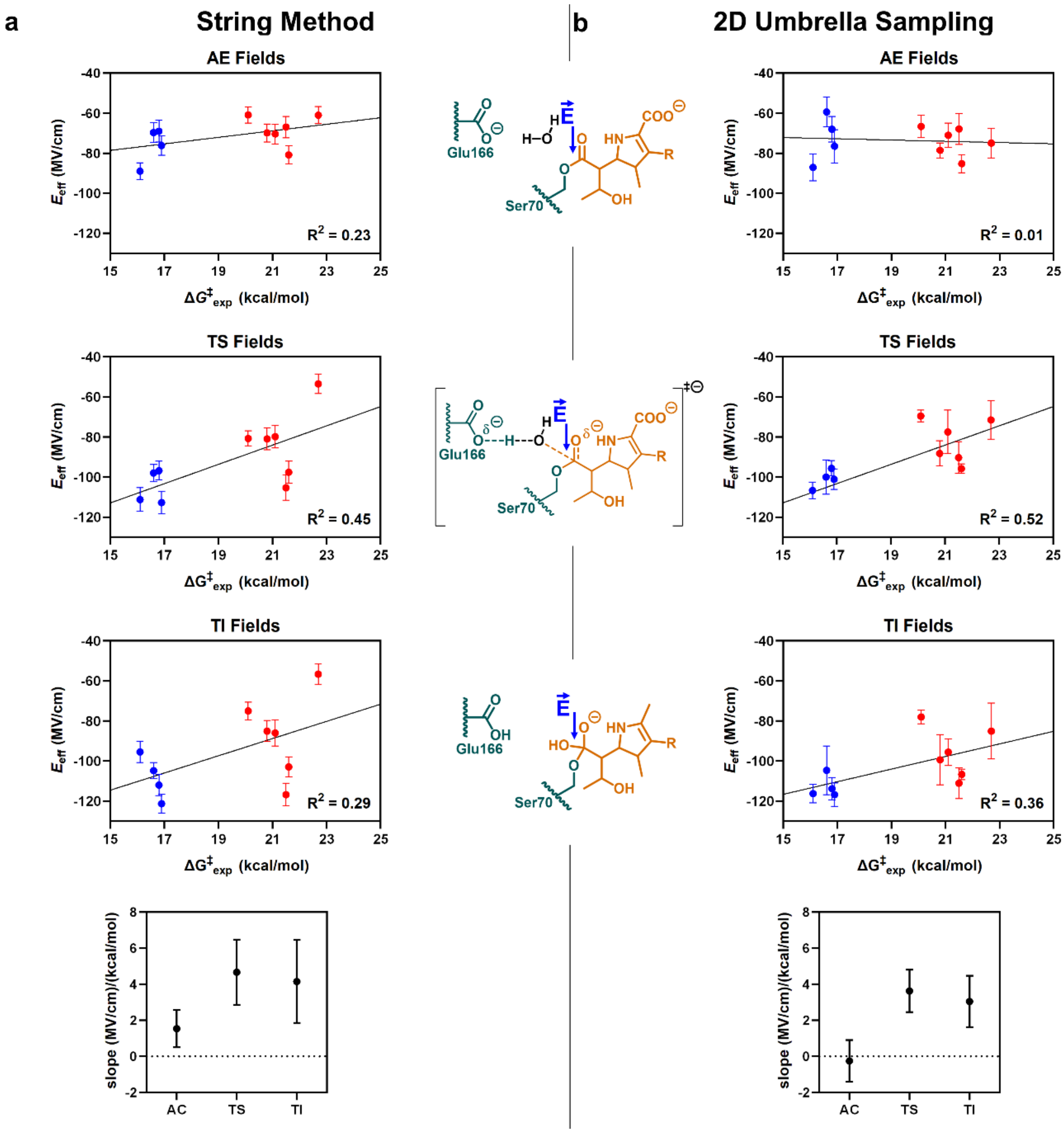

**Fig. S4 | Electric fields decomposition into individual states.**

**(a)** Total electric field in the acyl-enzyme (AE), the transition state (TS), and the tetrahedral intermediate (TI) calculated from trajectories based on the adaptive string method. The electric field varied with slopes of 1.6, 4.8, and 4.3 (MV/cm)/(kcal/mol) in the AE, TS, and TI states. **(b)** Analogous electric fields were calculated for three different states from trajectories based on 2D umbrella sampling. The electric fields varied with slopes -0.3, 3.7, and 3.1 (MV/cm)/(kcal/mol) in the AE, TS, and TI states.

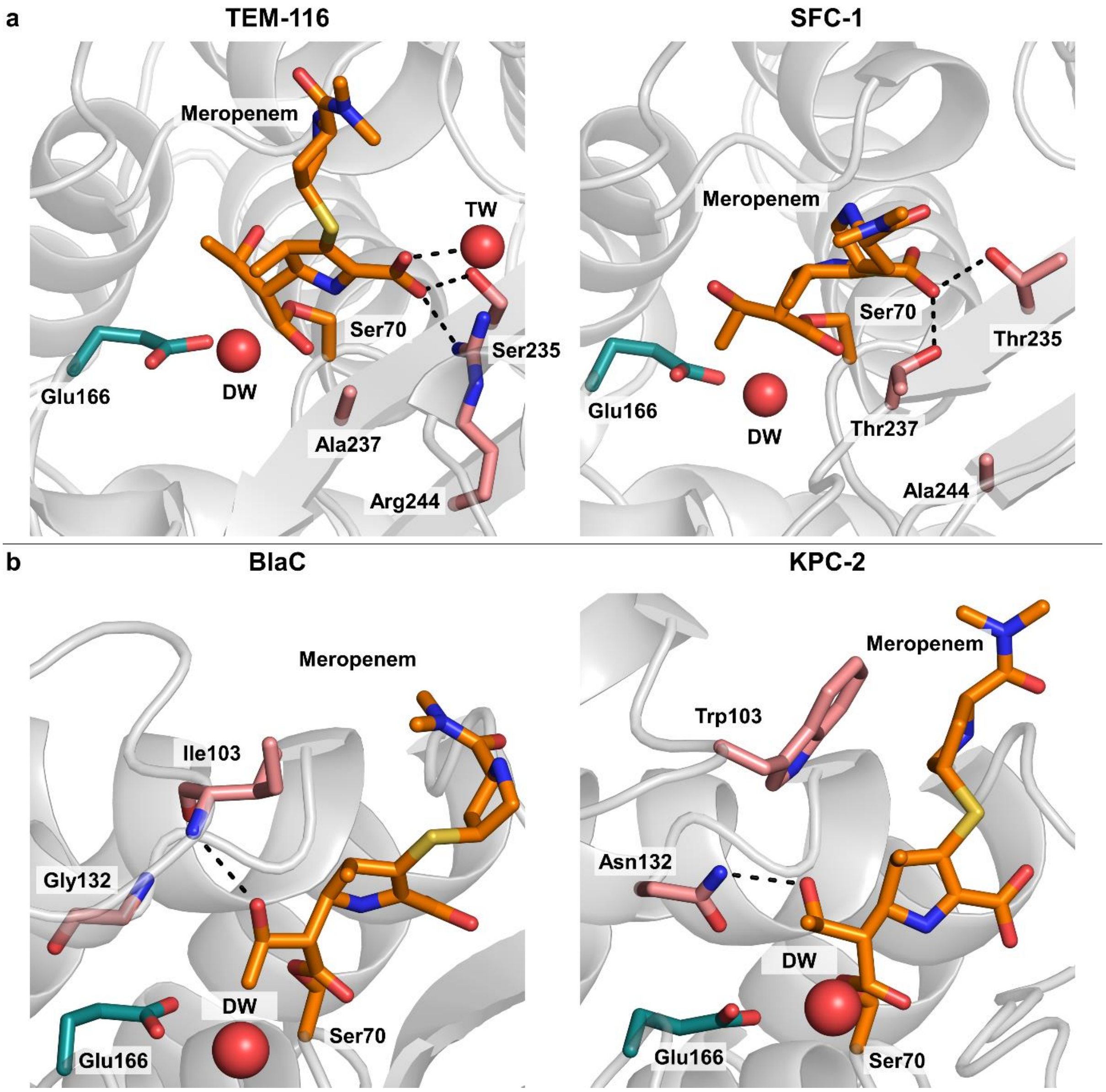

**Fig. S5 | Active site interactions in carbapenemases and non-carbapenemases.**

**(a)** Salt bridge interaction between Arg244 and the meropenem carboxylate in the active site of TEM-116 and SFC-1. **(b)** Structures of BlaC and KPC-2 showing how the Asn132Gly mutations changes the hydrogen bonding interaction with the 6-1R-hydroxyethyl group of meropenem, leading to formation of a new hydrogen bond with Ile103.

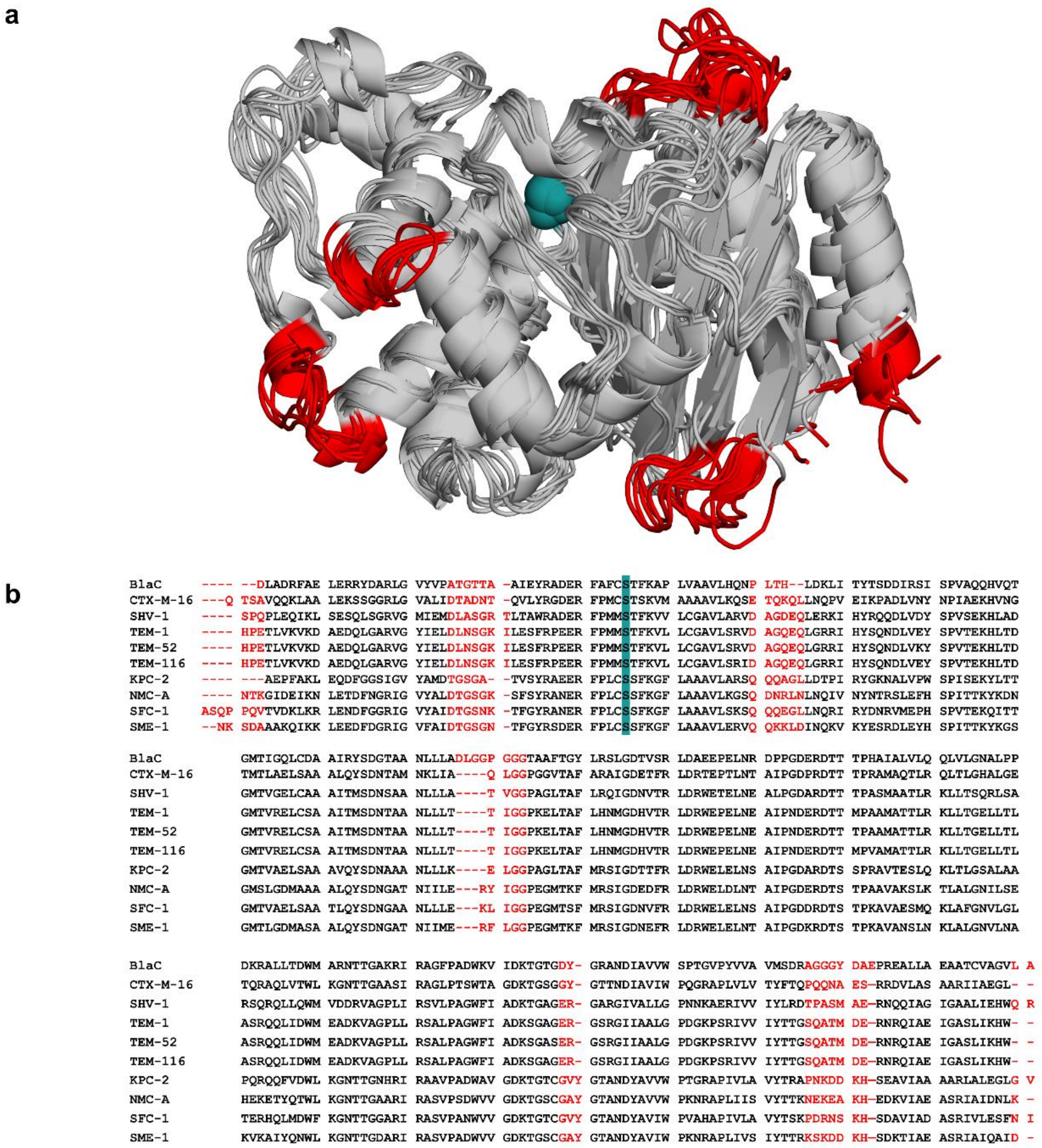

**Fig. S6 | Structure-based sequence alignment.**

**(a)** Cartoon showing loops excluded from the alignment and per-residue analysis due to varying lengths between variants (red). All loops are away from the active site (catalytic serine: teal spheres). **(b)** The excluded residues are highlighted in red in the multiple sequence alignment.

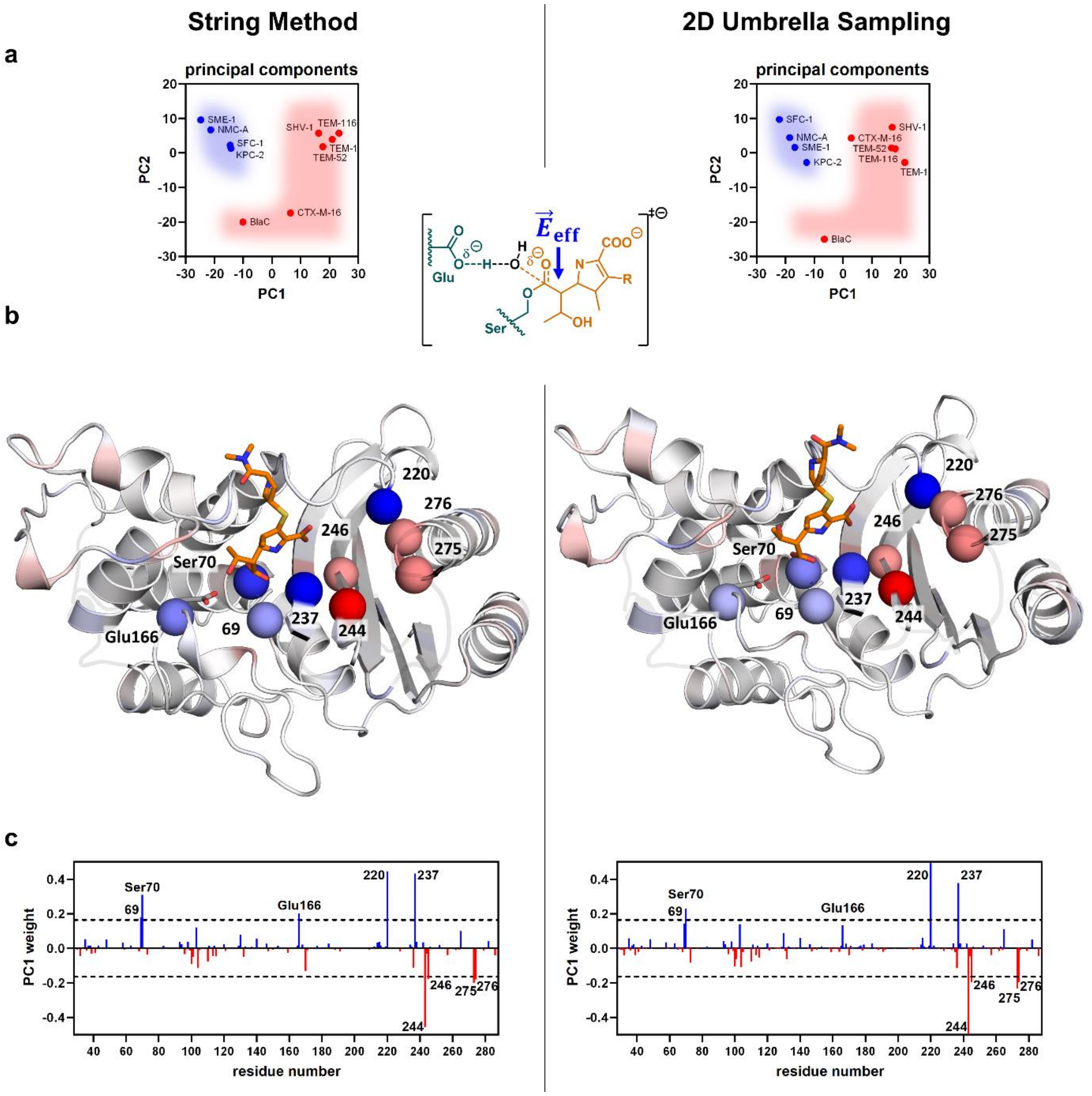

**Fig. S7 | PCA reveals residues key to electrostatic catalysis.**

Results from the string calculations **(left)** agree well with those from 2D umbrella sampling **(right)**. **(a)** PCA of the per-residue $E_{\text{eff}}$ clearly distinguishes carbapenemases (blue) from non-carbapenemases (red, shaded areas added for illustration only). **(b)** PC1 weights of each residue projected onto the β-lactamase scaffold reveals that a cluster of seven residues plus Ser70 and Glu166 (spheres) dominates $E_{\text{eff}}$ (blue: beneficial; red: detrimental, meropenem: orange sticks). **(c)** Electrostatic contribution of each protein residue suggested by PC1 weights. Key residues were identified as residues who's field varied by more than 2.5 standard deviation cutoff from all PC1 values (blue: beneficial, red: detrimental).
